# Age, experience, social goals, and engagement with research scientists may promote innovation in ecological restoration

**DOI:** 10.1101/2022.08.24.505135

**Authors:** Jakki J. Mohr, Tina M. Cummins, Theresa M. Floyd, Elizabeth Covelli Metcalf, Ragan M. Callaway, Cara R. Nelson

## Abstract

Innovation in ecological restoration is necessary in order to achieve the ambitious targets established in United Nations conventions and other global restoration initiatives. Innovation is also crucial for navigating uncertainties in repairing and restoring ecosystems, and thus practitioners often develop innovations at project design and implementation stages. However, innovation in ecological restoration can be hindered by many factors (e.g., time and budget constraints, project complexity, and others). Theory and research on innovation has been formally applied in many fields, yet explicit study of innovation in ecological restoration remains nascent. In order to assess the use of innovation in restoration projects, including its drivers and inhibitors, we conducted a social survey of restoration practitioners in the United States. Specifically, we assessed relationships between project-based innovation and traits of: the *individual practitioner* (including, for example, age, gender, experience); *company* (including, for example, company size and company’s inclusion of social goals); *project* (including, for example, complexity and uncertainty); and *project outcomes* (such as completing the project on time/on budget and personal satisfaction with the work). We found positive relationships between project-based innovation and practitioner traits (age, gender, experience, engagement with research scientists), one company trait (company’s inclusion of social goals in their portfolio), and project traits (project complexity and length). In contrast, two practitioner traits, risk aversion and the use of industry-specific information, were negatively related to project-based innovation. Satisfaction with work outcomes was positively correlated with project-based innovation. Collectively, the results provide insights into the drivers and inhibitors of innovation in restoration and suggest opportunities for research and application.

## Introduction

Innovation refers to a new idea, device, or method [1, 2] that may arise from novel recombination of existing ideas or the application of better solutions or unique approaches to solve existing problems [3]. Innovations can arise from many sources, including lessons learned from failed experiments, ideas from mavericks or “rogue” thinkers who inherently question everything, insights gleaned from adjacent endeavors, formal scientific research, and serendipity – the random “bolts of inspiration/insight” associated with genius inventors [4]. Similarly, innovations in ecological restoration can come from varied sources. For example, despite being controversial [5], challenging dogma and long-held assumptions about the way ecological restoration is conducted can stimulate new thinking [6]. Questions such as “should restoration always use native species?” [7] and “are local provenances always best?” [8] challenge assumptions in ways that can lead to innovation. Innovations in ecological restoration might also include the use of new technologies, such as drones or remote sensing that collect data more efficiently than labor-intensive techniques [9–12] and machine learning algorithms that improve decision making [13–15]. Innovations can arise from the use of new methods, such as novel approaches to growing seed stock [16] or leveraging genomics to address restoration concerns [17, 18]. Moreover, others advocate for bringing an entrepreneurial mindset to ecological restoration, such as regarding failure as a trigger for innovation [19; see also 5]. Indeed, “businesses [engaged in ecological restoration] are very well suited to fostering innovation in restoration,” and “the core values of many private firms can be aligned with innovation to support opportunistic tinkering” [20].

Innovation is crucial for both reversing high rates of environmental degradation that lead to the loss of biodiversity [21] and for achieving ambitious global targets for ecosystem restoration and repair (see the UN Decade on Ecosystem Restoration Report [22]. For instance, to meet commitments to restore millions of hectares of forests, innovation is identified as an urgent need for landscape-level planning and prioritization, seed sourcing and propagation, and monitoring success [23]. Hence, by bringing novel techniques and new approaches, innovation plays an important role in achieving goals for ecological restoration [6]. However, the factors that drive innovation in restoration are not well understood.

Despite its importance, innovation in restoration faces barriers and constraints. Resources for innovation are often absent or limited in ecological restoration compared to other industries such as agriculture, medicine, or business [6, 19]. Indeed, businesses must “navigate the many trade-offs and complexities in restoration projects” [24] to achieve ecological objectives while earning sufficient profit. Interestingly, despite its status as a dominant natural resource management activity, with over US$ 1 trillion spent annually in the global “restoration economy”25], research in restoration ecology has rarely addressed the perspective of the businesses engaged in the practice of restoration (see [24] for an exception). Yet, due to the inherent uncertainty of and possible failure from trying novel techniques, ecological practitioners can be reluctant to try new things. One practitioner summed it up:

> ...it is hard to do innovative things and still recognize the fact that down the road you’d be looked at as, was your project successful or not. In the end, that’s what people are interested in. They don’t really care if you use some innovative technology to get there or not” [24].

Thus, restoration practitioners who restore degraded landscapes face a paradox in innovation: on one hand, innovation is imperative to meet the global challenges and mandates for ecological restoration; on the other hand, innovation faces barriers and constraints. The purpose of our research was to understand the correlates (e.g., facilitators and barriers) of innovation in restoration projects. To our knowledge, this is the first study to examine this issue empirically in a restoration context.

We examined related fields for insights. In particular, business management and marketing have a long history of studying antecedents to innovation (e.g., [26, 27]). Indeed, Rogers’ seminal work on the diffusion of innovations focused on the area of agricultural innovations [28]. More recently, the broad field of environmental management has explored issues related to the uptake and adoption of new practices (e.g., [29–31]). Based on this prior research and adapting it to our restoration context, we grouped potential drivers and inhibitors into individual-level, company-level factors, and project-level traits. Individual traits such as age, gender, years of experience as well as risk tolerance (e.g., [29, 32]) are related to adoption of innovation; we explore whether and how these factors are related to innovation in the restoration context. Moreover, the adoption of innovations is related to information sources that an individual uses [31, 33]. Another variable related to innovation is the extent to which practitioners engage with research scientists [34]. Because the science/practice nexus is also critical in ecological restoration [35, 36], we explore both the extent to which a practitioner engaged with research scientists as well as the practitioner’s perceptions about the degree to which such engagement was helpful.

The relationship between company/organizational traits and innovation is also well-established in the literature [27, 31]. Although the business literature suggests that smaller companies tend to be more innovative and face fewer restrictions in using new ideas/techniques [37], the agricultural conservation literature suggest that size of an operation is positively related to use of innovation [31]. Hence, we examine the relationship between company size and innovation in the restoration context.

In business organizations, project complexity is negatively correlated with innovation [38]. In addition to project complexity, we also examined project length, as well as the degree to which project objectives were perceived as helpful (versus restrictive). Given the uncertain context in which restoration occurs, we also examined whether and how project uncertainty might affect the use of innovation in restoration projects. Because of the value of collaboration in stimulating innovation, other project-related variables included number of other businesses involved, the collaborative tenor of engagement with these other businesses.

Finally, people adopt innovations in order to achieve improved outcomes not attainable using existing methods, tools, and approaches. Because restoration outcomes may take years to unfold, we focused on the degree to which a particular project was completed on time and on budget. Moreover, we were curious about the relationship between innovation and the individual practitioner’s satisfaction with the work on the project.

In summary, our research addresses two primary questions: (1) what are restoration practitioners’ perceptions of the level of innovation in their projects? In other words, do they view their restoration work as relying on “tried-and-true techniques” compared to trying new methods and techniques? (2) How do (a) individual traits and personal characteristics, (b) company/ employer traits and characteristics, and (c) project traits and characteristics relate to respondents’ perceptions of the degree of innovation used in their projects? For example, do perceptions of project-based innovation vary by personal risk tolerance, company size, project uncertainty, project size, etc.? We also explored how project-based innovation related to outcomes, such as completing the project on-time/on-budget as well as the respondents’ personal satisfaction with the work. Our empirical findings provide insights for both research and practice regarding these issues.

## Methods

We focused on a wide variety of types of businesses engaged in ecological restoration in the United States, split roughly evenly between the scientific/engineering/design aspects of restoration and the physical construction/earth moving aspects [39, 40]. These firms work with agencies, non-governmental organizations (NGOs), and other stakeholders to execute restoration projects.

### Sampling frame and data collection

We designed and administered a questionnaire to restoration practitioners in the United States during the summer of 2017. Because the participant’s identity was not tied to the data and because risks to participants for filling out the questionnaire were minimal, the IRB approval fell under the “Exempt” category of review, meaning that an informed consent form was unnecessary (IRB #148-17). We obtained a database of 200 restoration practitioners from the Society of Ecological Restoration (SER) and, given our focus on the business perspective in restoration, we excluded email addresses that included a.gov or.edu extension. We used a single email solicitation, which included a letter from the SER executive director encouraging participation in our study. The email solicitation stated that participation was voluntary and that responses would remain anonymous. To ensure all respondents based their responses on the domain of ecological restoration, the survey instructions included the Society of Ecological Restoration’s definition of ecological restoration [41]. Respondents also were told they could skip any question and could quit the survey at any point. The fact that some participants skipped questions resulted in variation in sample sizes among analyses. A total of 97 surveys were completed (48.5% response rate).

### Measures

We developed measures for the constructs in our study based on studies of innovation in business [4, 42] and then contextualized these measures to the restoration context. Because they are psychometrically robust in capturing perceptions in survey research [43], we used Likert scales, which allow respondents to specify their level of agreement or disagreement to a series of statements (shown below). In addition to the measures for innovation, we also created measures to assess the possible barriers to and facilitators of innovation.

Measures were then pre-tested in individual sessions with three restoration practitioners who sat with the lead researcher as they filled out the survey. Each pre-test respondent spoke out loud about their reactions while completing the questionnaire. We made revisions based on their feedback—for example, a key insight offered was to measure the company’s inclusion of social goals in their restoration work—and the survey was piloted again by the same three practitioners and the research team before being coded into Qualtrics for electronic administration.

To assess innovation, respondents were prompted to select “The single project within the past three years that you are the most knowledgeable about” and then were asked to agree or disagree with three statements about project-based innovation:

- The project objectives were innovative.
- My company used new techniques on this project.
- My company introduced unproven methods on this project.

If an individual has not used a particular technique before, then that technique would be perceived as novel or innovative to that individual [26, 44]. In conjunction with these three items for Project-Based Innovation, respondents were asked to provide a brief description of the new technique or unproven method in the project. Measures for all other variables are presented in the supplementary information (S1 Appendix).

### Analysis

First, we relied on factor analysis to assess the dimensionality of our variables and for data reduction purposes for multi-item scales. Cronbach’s alpha provides a gauge of the internal consistency of multi-item variables. For scales of fewer than three items, coefficient alpha is not appropriate [45]; hence, we computed a Pearson correlation to assess the reliability of any two-item measures. Second, to assess the relationship between each of the sets of variables with the measures of innovation, our analysis relied on a correlational assessment. Although regression analysis would be a stronger test of the relationships between our predictor variables and innovation measures, our relatively low sample size, the option of skipping questions, and correlations between predictor variables (multicollinearity) precluded the use of regression.

## Results

This section first presents a description of the respondents’ traits, followed by the assessment of the measure’s reliability and dimensionality. We then present the results for the key variable, Project-Based Innovation, which is followed by the findings of individual, company, and project-level facilitators and inhibitors of innovation.

### Description of respondents

Table 1 provides a detailed breakdown of the respondents’ individual traits and characteristics (Panel A) as well as their company’s traits and characteristics (Panel B).

**Table 1.** Respondent profile. Panel A: Individual Characteristics

|  | **Mean/Std. Dev.** | **Median/Range** |
| --- | --- | --- |
| **Years in Restoration** | Mean = 19.76 years<br>Std. Dev. = 11.76 (n=85) | Median = 18.0 years<br>Range = 1-57 years |
| **Age (n=70)** | **n** | **%** |
| 20-30 years old | 5 | 7.1 |
| 31-40 | 13 | 18.6 |
| 41-50 | 20 | 28.6 |
| 51-60 | 20 | 28.6 |
| 60-70 | 11 | 15.7 |
| 71+ | 1 | 1.4 |
| **Gender (n=69)** |  |  |
| % Male/Female | 46/21<sup>a</sup> | 66.7%/30.3% |
| **Area of Expertise<sup>b</sup>** | **n** | **%<sup>c</sup>** |
| Ecological sciences | 59 | 22.4 |
| Plants / botany | 57 | 21.7 |
| Environmental sciences | 31 | 11.8 |
| Forestry and conservation | 25 | 9.5 |
| Water quality | 15 | 5.7 |
| Wildlife biology | 13 | 4.9 |
| Construction | 11 | 4.2 |
| River morphology / design | 7 | 2.7 |
| Civil engineering | 6 | 2.3 |
| Environmental engineering | 4 | 1.5 |
| Other 1<sup>d</sup> | 27 | 10.3 |
| Other 2 | 7 | 2.7 |
| Other 3 | 1 | 0.4 |

<sup>a</sup> n = 2 (3%) preferred not to answer; n = 21 did not answer/missing
<sup>b</sup> Area of Expertise exhibited no significant relationship with Project-Based Innovation
<sup>c</sup> Numbers sum to greater than 100% because respondents could select up to three areas of expertise; 23.3% of respondents identified 1 or 2 areas of expertise; 65.6% of respondents chose 3 areas; and 11.1% chose between 4-8 areas.
<sup>d</sup> Of the “Others,” the most common areas of expertise were: Architecture, Hydrology, Wetlands, and Horticulture

**Table 1.** Respondent profile. Panel B: Company Characteristics

|  | **Mean (Std. Dev.)** | **Median and Range<sup>a</sup>** |
| --- | --- | --- |
| Years working for current employer (n=84) | Mean = 11.06 years<br>Std. Dev. = 9.14 years | Median = 8.5 years<br>Range 1-42 |
| # of FTE at this office location (n=82) | Mean = 25.99 FTE employees<br>Std. Dev. = 35.44 | Median = 12<br>Range 1-150 |
| % of company's business that is in restoration (n=68) | Mean = 59.54%<br>Std. Dev. = 34.62 | Median = 57%<br>Range 1-100% |
| # of restoration projects per year (n=73) | Mean = 14.95<br>Std. Dev. = 21.34 | Median = 5<br>Range 1-100 |
| Average project size last three years (\$\$) (n=46) | Mean = \$102,521.74<br>Std. Dev. = \$135,698.15 | Median = \$50,000<br>Range 0-\$500,000 |
| This project's size (\$\$) (n=44) | Mean = \$516,081.80<br>Std. Dev. = \$819,279 | Median = \$280,000<br>Range = \$100-\$5,000,000 |
| **Geographic Region<sup>b</sup>** | **n** | **%** |
| Great Lakes Region | 19 | 23% |
| Northeastern Seaboard | 18 | 22% |
| Pacific Northwest | 12 | 15% |
| Southwest | 8 | 10% |
| Upper Rocky Mountains | 7 | 9% |
| California | 7 | 9% |
| Southeast | 6 | 7% |
| Great Plains | 4 | 5% |
| **Project Ecosystem<sup>b</sup>** | **n** | **% (of 90)** |
| Wetlands | 32 | 35.6 |
| Forest | 20 | 22.2 |
| Urban | 14 | 15.6 |
| Freshwater Aquatics | 14 | 15.6 |
| Coastal/Estuarine | 12 | 13.3 |
| Grasslands | 14 | 15.6 |
| Arid lands | 5 | 5.6 |
| Marine | 2 | 2.2 |
| Alpine/tundra | 2 | 2.2 |
| Other 1<sup>c</sup> | 13 | 14.4 |

<sup>a</sup> Outliers on some variables created skewness and hence, were removed from analysis: 3 outliers in # of FTE employees with 200, 350 and 800 FTE; 1 outlier in # of restoration projects per year with 200 projects; 3 outliers in Average project size (\$\$) with 700K, 1 mil, and 2 mil
<sup>b</sup> Neither Geographic Region nor Project Ecosystem exhibited significant relationships with Project-Based Innovation
<sup>c</sup> Other included Agricultural, Riparian and Floodplain

### Assessment of measures’ internal consistency and dimensionality

Because social science research relies on capturing human perceptions, measures are typically assessed for their internal consistency (reliability) as well as dimensionality. For example, multi-item measures (scales) are subjected to a factor analysis to assess whether the items load on one factor or if the scale is perhaps comprised of multiple factors (dimensions of the underlying construct). In addition, Cronbach’s alpha (greater than 0.7) is used to assess the measure’s internal consistency. We present all factor analyses in the Supporting Information (Tables S1-S7).

The items used to assess our key measure, *Project-Based Innovation,* all loaded on a single factor and exhibited a Cronbach’s alpha of 0.77, indicating acceptable reliability. The mean for Project-Based Innovation (on a five-point scale) was 3.02 (std. dev. = 0.80), with the distribution of responses shown in Figure 1. Notably, no respondents stated “Strongly Agree” in terms of their use of innovations on their projects.

**Fig 1.**
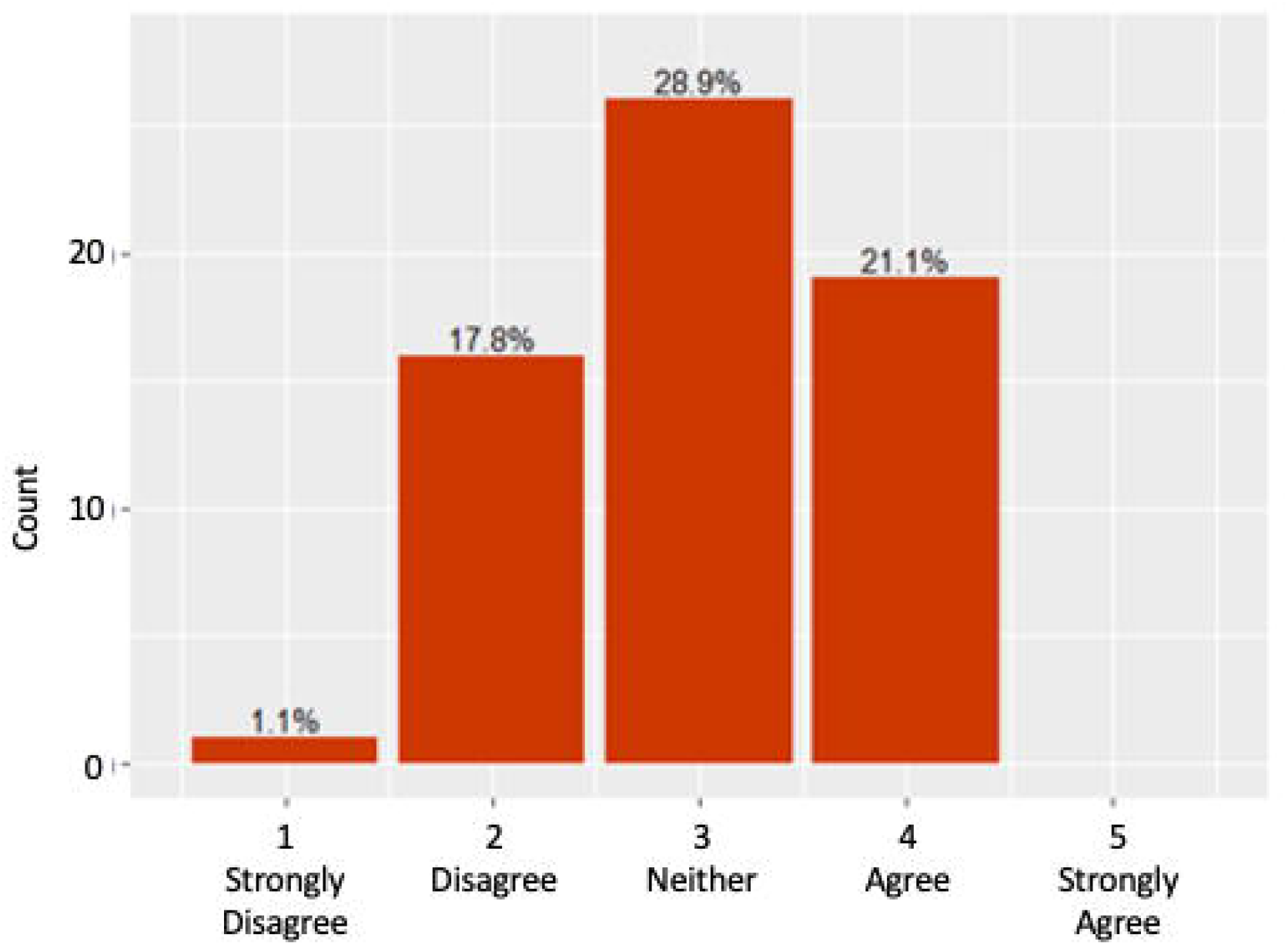
Distribution of Responses on Project-Based Innovation.

Respondents were asked to provide a brief description of the new technique or unproven method in the project (Table 2). In some cases, respondents stated that constraints in cost or design parameters forced them to innovate. In other cases, respondents developed innovations in response to dealing with invasive species or urban environments that required novel thinking. Still other respondents explained that their innovations applied “fairly standard practice” to a new region or ecosystem for which outcomes were uncertain. Other respondents identified using technologies such as drones for monitoring, new software for functional analysis, or the use of other new technologies for planning and monitoring restoration. One respondent had designed novel equipment for planting. In each case, responses indicated that the project required trying new methods or approaches for which evidence or guidelines were unavailable.

**Table 2.** Sampling of Qualitative Descriptions of Project-Based Innovations.

| **Description:** |
| --- |
| Used only salvaged material for erosion control and/or wetlands habitat |
| Creative in designing river channels (fish passages) due to cost |
| Lack of evidence of how to re-introduced 33 threatened and endangered plant and animal species |
| Collect seed from adjacent remnant properties in 30 mile radius |
| Used mulching techniques for invasive removal; new for our area but used in other geographic regions |
| Drones for monitoring |
| Innovative large-scale veg propagation to enhance ecological diversity at low cost |
| Creative stream baffles for erosion control and flood plain access |
| Lack of evidence on how to introduce native planting in urban environment to maximize diversity of wildlife habitats |
| Use pre-vegetated mats for trenching in perma-frost sites |
| Changed timing of invasive species control to be more effective |
| Wetland functional analysis as a design tool where it hadn't been used before |
| Re-use veg for erosion; |
| Recreate vernal pools for endangered species |
| Roller chipping site to reduce woody shrub coverage; not effective; returned to root raking after a few seasons |
| New application of old technique for erosion control on riverbank by using shrubs and woody debris to decrease alluvium flow into coral reefs |
| Phased invasive canopy removal to allow native canopy to adapt to new conditions |
| Proprietary/confidential |

### Facilitators and inhibitors of innovation

Here, we present the correlations between Project-Based Innovation and (a) the individual’s traits, (b) company characteristics, (c) project characteristics and (d) project outcomes. The correlational results report on the 66 participants for which we had complete data.

### Individual-level correlates of innovation

As Table 3 shows, perceptions of Project-Based Innovation were significantly correlated with age (r = 0.30, p<.01). Based on a median split on age (54.3% of respondents were younger than 50, while 45.7% of respondents were 51 or older), older respondents reported higher levels (mean = 3.17 and SD = 0.76) of Project-Based Innovation compared to younger respondents (mean = 2.90 and SD = 0.71). In addition, perceptions of Project-Based Innovation were significantly positively correlated with Years of Experience (r = 0.33, p<0.01). Males were significantly more likely to report that they engaged in Project-Based Innovation than females (r = 0.26, p <.05; mean = 3.18 and SD = 0.74 versus mean = 2.78 and SD = 0.71, respectively). A post-hoc t-test for gender was significant (t = 2.04, p<0.05). Reports of Project-Based Innovation also showed a significant negative correlation with Risk Aversion (r = −0.38, p<0.01); as Risk Aversion increased, Project-Based Innovation decreased.

**Table 3.**
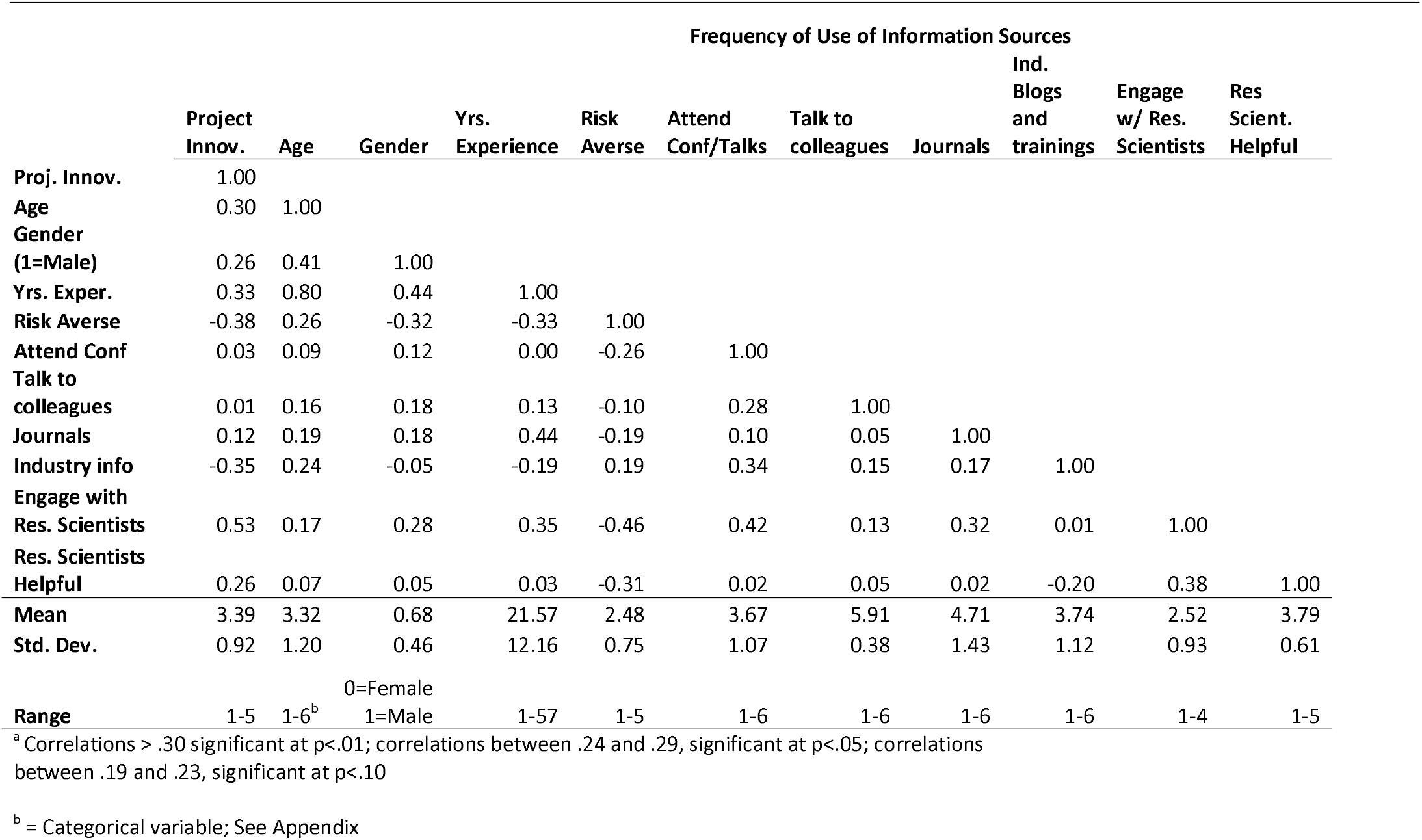
Correlational Analysis for Individual Traits and Characteristics^3^.

Regarding Sources of Information that respondents used, Project-Based Innovation was significantly negatively correlated with the use of industry trainings, blogs, and webinars (r = – 0.35, p<0.01); respondents who reported that they relied more heavily on these sources of information reported lower rates of Project-Based Innovation. No other information source factors (attending conferences; talking to colleagues; reading journals) exhibited significant correlations with Project-Based Innovation.

Respondents who engaged with research scientists—and perceived the engagement as helpful— reported higher rates of Project-Based Innovation (r = 0.53, p<0.01 and r = 0.26, p<0.05 respectively). An open-ended prompt to elaborate on the type of engagement with research scientists revealed that practitioners relied on such collaboration to assist in the design of experiments and data collection protocols, collection of specimens, molecular analyses, modeling assistance in estimates for ecological benefits and vegetation growth rates, designing assessment and monitoring plans, and accessing the “state of the science” in their work.

### Company-level correlates of innovation

As Table 4 shows, one company-level variable was significantly correlated with Project-Based Innovation: practitioners whose companies had a greater emphasis on social goals reported greater Project-Based Innovation than those whose portfolios had fewer or no projects with social goals (r = 0.25, p<0.05). Non-significant correlates of Project-Based Innovation were company size (FTE), the percent of the company’s business devoted to restoration, and the number of restoration projects per year.

**Table 4.** Correlational Analysis for Company Traits and Characteristics^a^.

|  | **Project**<br>**Innov.** | **Co Size**<br>**FTE** | **% Bus**<br>**Restoration** | **# Restor.**<br>**Projects** | **Social**<br>**Goals** |
| --- | --- | --- | --- | --- | --- |
| **Proj. Innov.** | 1.00 |  |  |  |  |
| **Co Size FTE** | 0.15 | 1.00 |  |  |  |
| **% Bus.**<br>**Restoration** | 0.10 | -0.17 | 1.00 |  |  |
| **# Restor.**<br>**Projects** | 0.03 | 0.18 | 0.31 | 1.00 |  |
| **Social Goals** | 0.25 | 0.13 | 0.12 | 0.09 | 1.00 |
| **Mean** | 3.39 | 25 | 58.32 | 15.08 | 2.91 |
| **Std. Dev.** | 0.92 | 31.60 | 32.49 | 21.64 | 0.96 |
| **Median** | 3.33 | 14.50 | 58.32 | 5.00 | 3.00 |
| **Range** | 1-5 | 1-135 | 1-100% | 1-100 | 1-5 |

<sup>a</sup> Correlations > .30 significant at p<.01 Correlations between .24 and .29, significant at p<.05 Correlations between .19 and .23, significant at p<.10

### Project-specific correlates of innovation

As Table 5 shows, two project-specific variables were significantly and positively correlated with Project-Based Innovation.

**Table 5.** Correlational Analysis for Project Characteristics^8^.

| | **Project**<br>**Innov.** | **Project Size**<br>**(\$\$)** | **Project**<br>**Size**<br>**(Relative)** | **Project**<br>**Complexity** | **Project**<br>**Length** | **# other**<br>**businesses** | **Collab.**<br>**Tenor** | **Obj.**<br>**Helpful** | **Climate-**<br>**Related**<br>**Uncertain** | **Company**<br>**Experience**<br>**Uncert.** | **Project**<br>**Politics**<br>**Uncert.** |
| --- | --- | --- | --- | --- | --- | --- | --- | --- | --- | --- | --- |
| **Project Innov.** | 1.00 |  |  |  |  |  |  |  |  |  |  |
| **Project Size \$\$** | 0.18 | 1.00 | | | | | | | | | |
| **Project Size**<br>**(Relative)** | 0.20 | 0.28 | 1.00 |  |  |  |  |  |  |  |  |
| **Project.**<br>**Complexity** | 0.27 | 0.27 | 0.51 | 1.00 |  |  |  |  |  |  |  |
| **Project Length** | 0.26 | 0.12 | 0.19 | 0.39 | 1.00 |  |  |  |  |  |  |
| **# other**<br>**businesses** | 0.21 | 0.05 | 0.22 | 0.20 | 0.19 | 1.00 |  |  |  |  |  |
| **Collaborative**<br>**Tenor** | 0.22 | 0.07 | 0.00 | 0.03 | -0.30 | -0.20 | 1.00 |  |  |  |  |
| **Helpful Obj.** | 0.20 | 0.15 | 0.18 | 0.04 | -0.12 | -0.02 | 0.06 | 1.00 |  |  |  |
| **Climate-related**<br>**Uncertainty** | 0.20 | -0.05 | 0.22 | 0.03 | 0.01 | 0.01 | 0.05 | -0.13 | 1.00 |  |  |
| **Company-**<br>**related**<br>**Uncertainty** | 0.16 | 0.14 | 0.08 | 0.29 | 0.32 | 0.05 | 0.09 | -0.13 | 0.08 | 1.00 |  |
| **Uncertainty due**<br>**to Project**<br>**Politics** | 0.17 | 0.15 | 0.20 | 0.31 | 0.16 | 0.27 | 0.02 | -0.27 | 0.22 | 0.17 | 1.00 |
| **Mean** | 3.39 | 516,081 | 2.38 | 2.42 | 4.28 | 5.80 | 3.34 | 4.23 | 2.70 | 1.96 | 2.11 |
| **Std. Dev.** | 0.92 | 666,360 | 0.73 | 0.55 | 4.20 | 6.34 | 0.35 | 0.72 | 0.79 | 0.73 | 0.94 |
| **Median** | 3.33 | 515,081 | 3 | 2.00 | 3.50 | 5.00 | 3.35 | 4.05 | 3.00 | 2.00 | 2.00 |
| **Range** | 1-5 | 100-5 Mill | 1-3 | 1-3 | <1-16 yrs | 1-50 | 1-5 | 1-5 | 1-4 | 1-4 | 1-4 |

<sup>a</sup> Correlations > .30 significant at p<.01; between .24 and .29, significant at p<.05

As relative project complexity and project length increased, so did Project-Based Innovation (r = 0.27 and r = 0.26, p<0.05 respectively). More specifically, more complicated projects exhibited a Project-Based Innovation score of 3.13 (std. dev. = 0.71) while easier projects exhibited a Project-Based Innovation score of 2.58 (std. dev. = 0.42). Projects shorter than 10 years exhibited lower levels of Project-Based Innovation (mean = 2.94, SD = 0.64) compared to projects longer than 10 years (mean = 3.15 and SD = 0.70). Five other project variables were marginally correlated with Project-Based Innovation: relative project size (r = 0.20), number of other businesses involved (r = 0.21), collaborative tenor of engagement with other businesses (r = 0.22), perceptions that objectives were helpful (r = 0.20), and uncertainty due to climate change (r = .20), all at p<.10. Uncertainty due to project politics was not significantly related to Project-Based Innovation.

### Outcomes from project-based innovation

As Table 6 shows, the correlation between Project-Based Innovation and Project Outcomes (completed on-time/on-budget) was non-significant. Respondents who reported higher levels of Project-Based Innovation reported higher Personal Satisfaction with their work (r = 0.42, p<0.05).

**Table 6.** Correlational Analysis for Project Outcomes’^1^.

|  | **Project**<br>**Innov.** | **Positive**<br>**Completion** | **Personal**<br>**Satisfaction** |
| --- | --- | --- | --- |
| **Proj. Innov.** | 1.00 |  |  |
| **Positive**<br>**Completion** | 0.05 | 1.00 |  |
| **Personal**<br>**Satisfaction** | 0.42 | 0.35 | 1.00 |
| **Mean** | 3.36 | 4.39 | 4.45 |
| **Std. Dev.** | 1.02 | 0.65 | 0.91 |
| **Range** | 1-5 | 1-5 | 1-5 |

<sup>a</sup> Due to lower sample size (n=22 respondents for completed projects), correlations from .42 - .49, significant at p<.05; correlations between .36 - .41, significant at p<.10

## Discussion

The purpose of our research was to understand the correlates (e.g., facilitators and barriers) of innovation in restoration projects. Given the critical importance of innovation to restoration, understanding practitioner perspectives on innovation and the traits associated with innovative practices are greatly needed. Our study is the first to investigate practitioner perspectives on innovation in ecological restoration. This topic is of vital importance: although innovation is often needed to achieve successful restoration, innovation is inherently risky. The trade-off between sticking with a tried-and-true approach versus trying something new–and perhaps risky but with a potentially higher payoff in terms of achieving successful outcomes—is one that warrants attention.

Practitioners rated their Project-Based Innovations as about average, with no respondents expressing strong agreement about the innovativeness of their project objectives, techniques and methods. Those who reported higher Project-Based Innovations were older, more experienced, men who were less risk averse; they engaged with research scientists and perceived such engagement as being helpful; their companies’ projects included a focus on social goals; and more complex and longer projects were correlated with higher levels of Project-Based innovation. Here, we situate these findings relative to related literature, acknowledge our study’s limitations, and offer suggestions for future research.

The finding that older respondents reported higher levels of Project-Based Innovation is inconsistent with theory for the adoption of innovation, which posits that adopters of new innovations are generally younger [26]; see also [31]’s review which found that older farmers are less likely to adopt agricultural conservation practices. Perhaps in the field of restoration, age engenders greater confidence, giving older practitioners the necessary perspective to experiment with new techniques and novel methods. This possible explanation is supported by our finding that more experienced practitioners reported higher levels of Project-Based Innovation. (Recall from Table 3 that age and experience were highly correlated (r = 0.80).

On the other hand, consistent with innovation theory [26], our results show that males tend to report higher levels of Project-Based Innovation than females. Given that in our study, women skewed younger than men, and because gender is significantly correlated with experience in our study (with men tending to have more experience than women), it is difficult to get a clear picture of the relationship between Project-Based Innovation and gender relative to age and experience. The interplay between these variables would be an interesting area of research, particularly in light of [31], that found women were more likely than men to adopt novel agricultural conservation practices.

The strong negative relationship between Risk Aversion and Project-Based Innovations is consistent with prior research showing that people who are risk averse are less likely to innovate [32]. Consistent with [24]’s recommendations based on qualitative findings, our study reinforces the value of encouraging restoration practitioners to be aware of their own risk aversion and to balance it with a willingness to try new things.

The negative relationship between the frequency of usage of industry trainings, blogs, and webinars as sources of information and Project-Based Innovation is intriguing. Typically, adopters of new innovations tend to rely more extensively on industry sources of information [26]. In contrast, our research shows that in the field of ecological restoration, practitioners who relied on these sources of industry-based information were less likely to engage in Project-Based Innovation. Perhaps industry-based information reinforces existing industry practices rather than encouraging novel techniques. Teasing out this relationship would be a fruitful area for additional inquiry.

In contrast to the negative relationships between use of industry-related information and Project-Based Innovation, practitioners who engaged more extensively with research scientists, and who found such engagements helpful and valuable, reported greater Project-Based Innovation. Although scientists are sometimes criticized for being out of touch with “on-the-ground” realities of practitioner needs [24, 46], our findings suggest such collaboration can offer an important source of new insights and methods. Exploring venues and opportunities to facilitate such engagement would be valuable to sparking innovation in ecological restoration.

Our finding that a company’s inclusion of social goals in their work is positively related to Project-Based Innovation suggests that broadening the scope of restoration projects beyond ecological domains can stimulate novel techniques and thinking. The underlying principles of ecological restoration state the importance of stakeholder engagement and social goals [47]. The specific mechanisms by which social goals engender innovation warrants further investigation. Perhaps social goals provide a new lens through which restoration practices are viewed, leading to questioning standard practice and lending new insights. Or, perhaps the inclusion of social goals provides greater engagement with other stakeholders, in turn generating opportunities for co-discovery and co-innovation of on-the-ground techniques [48]. Or possibly restoration ecologists who are open to social goals may also be open to innovation.

Prior studies of innovation in business suggest that complexity can inhibit innovation [38] and longer projects may have greater unknowns. However, our study found positive relationships between both project complexity and project length with Project-Based Innovation. Perhaps longer timelines allow, or even facilitate, more experimental thinking. Similarly, perhaps more complex projects spark more out-of-the-box thinking. Looking forward, awareness of project complexity and longer time frames may stimulate innovation and creative thinking to emerge from restoration practitioners.

Finally, our finding that an individual’s personal satisfaction with their work on the project is positively related to Project-Based Innovation suggests that using novel techniques and new methods offers a sense of gratification and fulfillment. (Given our cross-sectional data, it is also possible that someone who really enjoys their work might be more likely to innovate.) This finding is particularly noteworthy, given that Project-Based Innovation was unrelated to whether the project was completed on time or on budget. In other words, even if a project’s outcomes were not positive, individuals experienced greater sense of satisfaction with more innovative projects. Practitioners should consider how to best allow for innovation where staff feel safe to experiment and are rewarded for innovative thinking. The finding that Project-Based Innovation was not correlated with project outcomes is perhaps not surprising, given that some innovations are likely to contribute to improved outcomes while other innovations are likely to not work out as expected.

The large number of non-significant correlations is consistent with [31], whose review of 93 studies revealed that few independent variables have a consistent relationship with farmer adoption of conservation practices. Likewise, the non-significant relationships in our sample could be a function of the high variance in our sample, for example, the wide variety of backgrounds, training and the types of ecosystems in which the restoration was conducted. We expected that by relying on individual respondents’ perceptions of innovation (cf. [49]), our measures would be able to handle such nuances and subtleties. However, various types of innovations—whether new products or technologies (i.e., new type of sensor), new procedures or methods (e.g., genomics or algorithms), or even new approaches (public/private partnerships)—might exhibit different correlates. Additionally, different types of restoration practitioners (e.g., wildlife managers vs. civil engineers) likely experience different facilitators of and barriers to innovation. Similarly, restoration on government contracts might exhibit different triggers of and propensities for innovation than restoration of private lands. Whether our U.S.-based findings might differ from other geographic regions is also an open question. Understanding these and other contextual nuances offers the potential for additional insights into facilitators and barriers of innovation in ecological restoration.

We relied on a well-established approach to measuring perceptions in social science research: Likert scales. It would be interesting to consider other ways to assess innovation in this context as well, for example through measures of expert assessment of the novelty of particular designs and practices. Moreover, we encourage future research on specific restoration innovations other than the project-based innovations our measures focused on. For example, social innovations, such as public and private partnerships or processes for engagement and co-innovation, may be as important as ecological innovations in coping with challenges in ecological restoration.

Another approach to studying the use of innovations in ecological restoration would be to consider specific messaging or communications strategies that might be used proactively to stimulate adoption of innovation. For example, [31] suggested the importance of tailoring different messages to different types of farmers to stimulate adoption of conservation measures. Although our study examined sources of information that practitioners rely on, we did not examine the content or strategies those sources of information conveyed. Specific communication strategies, such as those used to facilitate comparisons to others’ behaviors, can be used to “nudge” people along the path to adoption [50, 51], a potentially useful approach to upscaling innovation in ecological restoration. Related to communication strategies, social networks are an important aspect of influence and information (e.g., [29]). Given that different types of network relationships exert different levels of influence on individuals’ behavior, it would be helpful to explore the impact of individuals’ social connections on their use of innovation in ecological restoration.

An issue related to individual practitioner propensity to use innovation is the diffusion of innovation across a discipline, sometimes referred to as “scale up” or uptake. Based primarily on adoption and diffusion of innovation [26], research in the related discipline of conservation has used both case studies [30] as well as quantitative modeling (Mills et al. 2019) to explore how innovations diffuse throughout a particular discipline (see also [53, 54]).

Finally, the relationship between “best practices” in a discipline and innovation offers the potential for valuable insights [55]. Best practices are often codified in industry standards regarding how to perform a particular task or function. The role of such standards, particularly in young fields, may lead to early reification of practices and techniques that might stymie novel thinking [56]. Finding the sweet spot between promoting standards that improve the practice of ecological restoration [47, 57], while at the same time encouraging innovation, would offer fruitful results in ecological restoration.

## Conclusion

In summary, our findings uncovered a number of factors that may facilitate or inhibit project-based innovation in ecological restoration. Older, more experienced practitioners are more likely to report the use of innovations in their restoration projects, while those with greater risk aversion are less likely to report innovative solutions in their work. In addition, individuals who engage with research scientists, and who find those engagements helpful, report greater use of innovative solutions, while those who access industry information report less innovative solutions. Those who work for companies that include social goals report the use of innovations in their restoration projects, as do those who work on more complex, longer projects. Finally, Project-Based innovation is related to satisfaction with the work, indicating either that innovation results in greater satisfaction or that satisfied individuals are more open to innovation. Regardless, innovation appears to be a meaningful aspect of an individual employee’s work. Based on these findings, we offer the following recommendations to improve innovation in ecological restoration:

- To stimulate innovative practices in early-career practitioners, collaboration between older and younger practitioners may be beneficial.
- To increase rates of innovation, practitioners should be aware of their own risk aversion and attempt to balance it with a willingness to try new things; their companies may also consider ways to decrease the negative consequences of risk for their employees.
- To generate new techniques and practices, engagement with research scientists and inclusion of social goals in ecological restoration can offer favorable opportunities.
- Longer and more complex projects may increase opportunities for innovation.

Our hope is that these insights encourage practitioners to understand their own aptitude for innovation and to explore new approaches to ecological restoration in order to meet the challenges of the coming Decade of Ecological Restoration [58, 59].

## Acknowledgements

Thank you to the Society for Ecological Restoration for allowing us access to its database for our survey and for sending along a letter to encourage participation in the study. The first author thanks feedback from the University of Queensland Centre for Biodiversity and Conservation Science (CBCS) and ARC Centre for Excellence in Environmental Decisions (CEED) for helpful comments, and Bethanie Walder for invitations to present this research at SER conferences as well as for her helpful comments on survey design and interpretation of results. The first author thanks John DeArment and Pedro Marques for their time and thoughtful responses during the pre-tests. She also thanks Jessica Diehl for her capable assistance in building the sampling frame, programming the survey, and collecting the data.

## Supporting information

**S1 Appendix. Measures for Individual, Company, and Project-Level Variables**

**Table S1. Factor Analysis for Innovation Items**

**Table S2. Factor Analysis for Risk Aversion**

**Table S3. Factor Analysis for Sources of Information Items**

**Table S4. Factor Analysis for Perceptions of Engagement with Research Scientists**

**Table S5. Factor Analysis for Items for Collaborative Tenor of Relationships**

**Table S6. Factor Analysis for Level of Uncertainty Introduced**

**Table S7. Factor Analysis for Project Outcome**

## S1 Appendix. Measures for Individual, Company, and Project-Level Variables

**Project-Based Innovation:** Three items anchored on a five-point Strongly Disagree to Strongly Agree scale; loaded on one factor.

*My company used new techniques on this project.*

*My company introduced unproven methods on this project.*

*Project objectives were innovative.*

### Individual Traits

Age: 6 categories (20-20; 31-40; 41-50; 51-60; 61-70; 71+)

Gender: male, female, prefer not to answer

Area of Expertise: Ecological science, plants/botany, environmental sciences, forestry/conservation, water quality, wildlife biology, construction, river morphology/design, civil engineering, environmental engineering, other

Risk Aversion: three items anchored on a five-point Strongly Disagree to Strongly Agree scale; loaded on one factor.

- I take minimal risks to avoid potential negative outcomes.
- I try to avoid risks at all costs.
- It is not worth it to take substantial risks just for the hope of achieving a positive outcome.

Information Sources: Frequency of use of six information sources (anchored on a six-point scale ranging from 1=Never; 2 = Less frequently than one every two years; 3 = Once every two years; 4 = Once a year; 5 = Twice a year; to 6 = Once a quarter or more frequently). Loaded on three factors.

- Professional conferences in my field
- Academic or university sponsored talks/presentations
- Practitioner journals and magazines
- Academic journals like Restoration Ecology
- Online webinars/trainings
- Industry blogs and online resources
- Trainings by private companies such as the Wetlands Institute or the Rosgen Center
- Talking to other colleagues and professionals in the discipline (did not load on any factor)

Extent of Engagement with Research Scientists. To what extent have you engaged with research scientists (non-practitioners) in the course of doing your ecological restoration work over the past three years (1 = Not at all; 2 = A minimal extent; 3 = A moderate extent; 4 = A great extent).

Perceptions of Helpfulness of Engagement with Research Scientists. (For those who said “A moderate” or “A great” extent to engagement above, they were asked to respond to the following five items on a five-point scale; loaded on one factor:

- helpful
- valuable
- realistic
- spot-on
- cutting-edge

### Company Variables

Years working for current employer

\# FTE at respondent’s office location

% of company’s business in restoration

Average # of restoration projects/year (over past three years)

Extent of Social Goals in company work in past three years (four-point scale 1 = Not at all; 2 = A minimal extent; 3 = A moderate extent; 4 = A great extent): “Some ecological restoration projects have goals oriented towards society as a whole and they might include things like public use and recreation, economic livelihoods from the land, and so on. These are sometimes referred to as “social goals.” In the past three years, to what extent have the projects your company worked on had social goals (e.g., public use and recreation) as part of the purpose of the ecological restoration?”

**Project Variables** Respondents were prompted: “For the following questions, please select the single project within that past three years that you are the most knowledgeable about.”

Project size ($ value)

Relative project size (1 = smaller than usual/ 2= typical/ 3 = larger than usual)

Project complexity (1 = easier than usual/ 2 = typical/ 3 = more complex than usual)

Project length/duration (years)

\# of other businesses involved in project (and # of other businesses previously worked with and

% of other businesses previously worked with, e.g., # previously worked with/#involved) Tenor of collaboration with those businesses (4 items: anchored on a five-point Strongly Disagree to Strongly Agree scale); loaded on one factor.

- In general, the various businesses worked well together.
- The parties effectively worked through differences.
- Communication between the various parties was effective.
- The various businesses seemed to step on each other’s toes. (Reverse scored)

Project Objectives (2 items: anchored on a five-point Strongly Disagree to Strongly Agree scale):

- They allowed high quality restoration.
- They were realistic.

Project Uncertainty: Level of uncertainty introduced by the following six dimensions; loaded on three factors.

- The vagaries of nature (floods, weather, drought, fire).
- Climate change induced uncertainties.
- Experience of my company with this type of project.
- Restoration methods/protocols for this project domain.
- Disagreements between/among various stakeholders /businesses/ organizations/citizen collaboratives, etc.
- Politics external to the project.

**Project outcomes**: For respondents who stated that the company’s work on the project had been completed (n=25); loaded on one factor. To what extent was the project:

- completed on time
- completed on budget
- accomplished the design goals
- met major milestones as planned

“How satisfied are you, personally, with your own work on this project?” (Extremely Dissatisfied to Extremely Satisfied)

## Notes

### Competing Interest Statement

The authors have declared no competing interest.

### Summary of Updates

Includes the Appendix with the items used in the questionnaire.

