## Supplemental files for "Age, experience, social goals, and engagement with research scientists may promote innovation in ecological restoration"

“How satisfied are you, personally, with your own work on this project?” (Extremely Dissatisfied to Extremely Satisfied)

**Table S1. Factor Analysis for Innovation Items^a^**

“Project Innov.”

My company used new techniques
 on this project	………….	.88
My company introduced
 unproven methods on this project …..	.67

Project objectives were innovative…….	.53

Variance explained	32%
Mean (Std. Dev.)	3.39 (0.92)
Coefficient Alpha	.77

^a^n=66 observations, including 7 mean substitutions

**Table S2. Factor Analysis for Risk Aversion^a^**

Factor

I take minimal risks to avoid potential negative outcomes.	.73

I try to avoid risks at all costs.	.71

It is not worth it to take substantial risks just for the hope

of achieving a positive outcome.	.64

Variance Explained	48%

Mean (Std. Dev.)	2.48 (.75)

Coefficient Alpha	.72

**^a^** 75 observations, no mean substitutions

**Table S3. Factor Analysis for Sources of Information Items^a^**

F1	F2	F3
Journals	Conferences/	Industry
Presentations	Training

Professional conferences in my field	.59

Academic or university sponsored talks/presentations	.78

Practitioner journals and magazines	.71

Academic journals like Restoration Ecology	.96

Online webinars/trainings	.36

Industry blogs and online resources	.82

Trainings by private companies	.35

such as the Wetlands Institute or the Rosgen Center

Talking to other colleagues and professionals in the discipline**

Variance Explained	21%	16%	14%
Mean (Std. Dev.)	4.71(1.43)	3.67(1.07)	3.74(1.12)

Pearson Correlation	r=.69, p<.000	r=.45, p<.000	α= .52

**^a^** 66 observations, no mean substitutions

^b^ Did not load on any factor and hence, examined as a separate item.

**Table S4. Factor Analysis for Perceptions of Engagement with Research Scientists^a^**

Factor:

“Helpful”

Helpful	.72

Cutting Edge	.75

Realistic	.71

Spot-On	.80

Valuable	.64

Variance Explained	53%

Mean (Std. Dev.)	3.8(.66)

Coefficient Alpha	.85

^a^ In general, how would you characterize your engagement with these research scientists?  
 Please answer how strongly you agree or disagree with the characteristics.

**Table S5. Factor Analysis for Items for Collaborative Tenor of Relationships^a^**

Factor

In general, the various businesses worked well together.	.71

The parties effectively worked through differences.	.54

Communication between the various parties was effective.	.93

The various businesses seemed to step on each other’s toes. (R)	.68

Variance Explained	53%

Mean (Std. Dev.)	4.13(.64)

Coefficient Alpha	.80

**^a^** 66 observations, 15 mean substitutions

(R) means reverse-scored

**Table S6. Factor Analysis for Level of Uncertainty Introduced By:^a^**

F1	F2	F3

“Project Politics”	“Company”	“Climate”

The vagaries of nature	.37

(floods, weather, drought, fire).

Climate change induced uncertainties.	.92

Experience of my company

with this type of project.	.86

Restoration methods/protocols

for this project domain.	.66

Disagreements between/among	.82

various stakeholders /businesses/

organizations/citizen collaboratives, etc.

Politics external to the project.	.73

Variance Explained	21%	21%	18%

Mean (Std. Dev.)	2.11(.94)	1.96 (.73)	2.7 (.79)

Pearson Correlation	r = .62, p<.000	.56, p<.000	.36, p<.002

**^a^** 66 observations, 11 mean substitutions

**Table S7. Factor Analysis for Project Outcomes^a^**

Extent to which the project was:	Factor

Completed on time	.88

Completed on budget	.85

Accomplished the design goals	.39

Met major milestones as planned	.78

Variance Explained	57%

Mean (Std. Dev.)	4.4(.65)

Coefficient Alpha	.82

**^a^** For respondents who stated that the company’s work on the project had been completed (n=22)
